# Flexible visual learning and memory in nectar foraging hornets

**DOI:** 10.1101/2023.02.06.527296

**Authors:** Mathilde Lacombrade, Monica Doblas Bajo, Naïs Rocher, Emmanuel Navarro, Christian Lubat, Fanny Vogelweith, Denis Thiéry, Mathieu Lihoreau

## Abstract

Pollinators, such as bees, develop flexible memories of colours, patterns and shapes, for efficient flower recognition. Here we tested whether other insects facing the same foraging problem have evolved similar cognitive abilities for flexible learning. We trained wild hornets from two species commonly found in Europe (*Vespa velutina nigrithorax* and *Vespa crabro*) to associate sucrose solution rewards to colour stimuli in a Y maze. Hornets of both species succeeded in differential and reversal learning, and developed short and long-term memories of the learnt associations. Thus, just like bees, hornet foragers can learn various visual cue-reward associations and remember them over several hours or days for selecting flowers. Our study in non-model species illustrates how standard conditioning approaches can be used to explore and compare the cognitive abilities of animals with similar foraging ecology.

## Introduction

Nectar foraging animals, such as bees, butterflies, birds and bats, have evolved a rich cognitive repertoire for flower recognition [1–4]. Bees, for instance, develop accurate visual memories of shapes, colours, and patterns to exploit best rewarding flowers [5–10], and some of these information can last in memories for days or weeks [11], allowing for flower specialization (i.e. flower constancy [12]). At the most basic level, foragers can discriminate flowers by learning associations between visual cues and a reward or the absence of it (differential learning [13]). Importantly, they can also exhibit some cognitive flexibility in order to update these learnt associations (reversal learning [14]) or learn new ones as the rewarding value of flowers change through time, for instance if learnt species become unavailable and new ones start blooming. Consequently, we would expect such flexible visual associative learning and memory to be broadly observed across nectar foraging species.

Wasps constitute a large group of Hymenopteran insects, phylogenetically close to bees, and in which many species frequently forage floral nectar [15] (although their main source of food is protein [16]). These insects are well-known to use visual cues for place learning [17] and nestmate recognition [18]. Recent studies using appetitive conditioning also reported their ability to learn pictures of human faces [19], patterns [20] and colours [21]. However, little is known about the ability of wasp foragers to be flexible in these visual learning, in order to update preferences with changes in resource quality, and keep these information in memory for optimizing nectar foraging trips, just like bees do [22].

Here we investigated visual learning and memory in the two main hornet species found in Europe: The European hornet (*Vespa crabro*) and the invasive Yellow-legged hornet (*Vespa velutina nigrithorax). V. crabro* is present in Europe since at least two centuries [23] while *V. velutina* was first recorded in 2004 [24]. Since these social hornets occasionally forage on flowers, we hypothesized that they should exhibit flexible visual learning. We tested this hypothesis by adapting visual appetitive conditioning protocols previously developed for bees [25], using a semi-automatic Y-maze.

## Methods

### Hornets

We caught wild nests of *V. crabro* (n=1) and *V. velutina* (n=3) during autumn 2022 (see Table S1) and cooled them (4°C) for 24h. We paint marked a subset of adults in each nest with a unique color code on their thorax for individual identification and transferred the nests into plastic boxes (24 cm h x 32 cm l x 32 cm d – Figure 1A). We kept the nests in an experimental room at ambient temperature (20-22°C) and provided hornets with *ad libitum s*frozen honeybees (source of proteins) and 40% (v/v) sucrose solution directly into the box.

**Figure 1:**
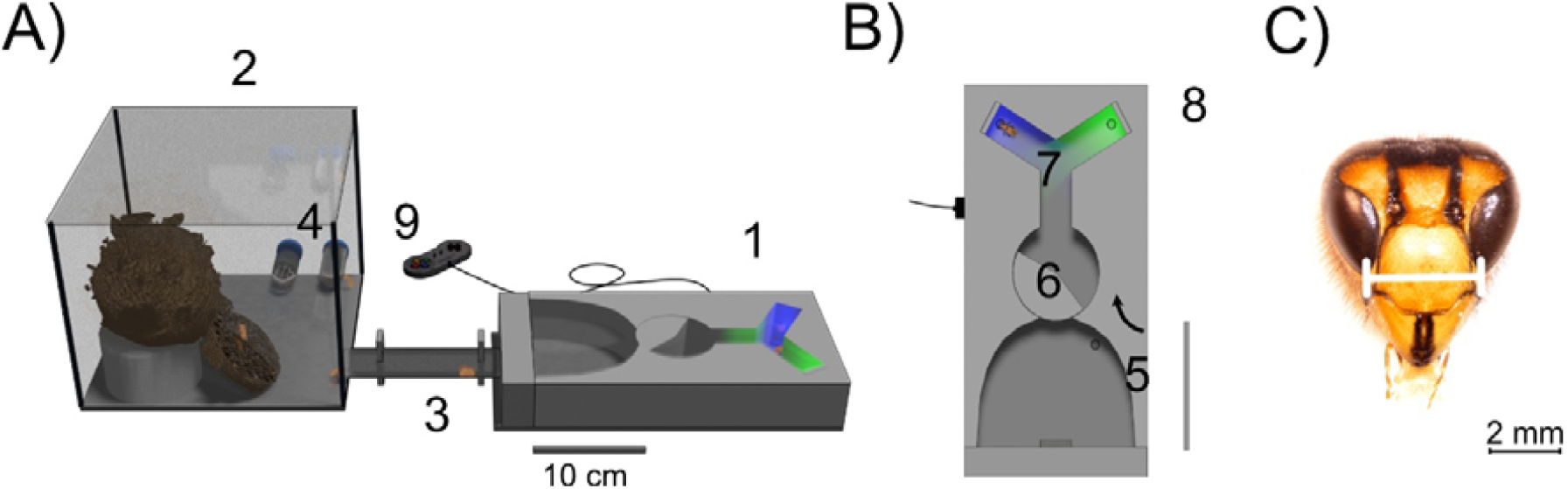
A) Overall view of the setup. (1) Y maze connected to (2) nest box through a (3) tunnel with shutters. (4) Feeders containing *ad libitum* food (honeybees and sucrose solution). B) Detailed view. The Y maze was dung in Styrofoam and covered with transparent plexiglass. (5) Pre-training feeder. (6) Turnstile entrance/exit door. (7) Feeding holes containing sucrose solution (positive reinforcement) or water (negative reinforcement) associated with (8) LED light displayed on the back wall (visual stimuli). (9) Turnstile and LEDs (on/off, change colours) were remote controlled using a manual controller. C) Head width measurement (white line).

### Y maze

We connected each nest box to a homemade Y maze using a clear transparent plastic tunnel (52 cm l, 2 cm Ø) with shutters to control the flux of foragers (see details in Figure 1B). Hornets could enter and exit the Y maze through a turnstile door at the entrance. The first branch led to two identical arms, each containing a feeding hole (0.2 ml) at its extremity. Depending on the protocol, the hole could contain sucrose solution, water, or nothing. The back wall of each arm was illuminated by colored LEDs through a diffuser (blue light: λ = 465-467 nm, intensity = 180-200 mcd; green light: λ = 522-525 nm, intensity = 660-720 mcd). Activation of the turnstile door and the LEDs were remote controlled by the experimenter.

### Pre-training

We pre-trained hornets to collect 40% (v/v) sucrose solution *ad libitum* on a feeder placed at the entrance of the Y maze (Figure 1B). The feeder was a lidless transparent 0.2 ml Eppendorf inserted into the floor of the maze. We considered all hornets that made at least 3 visits to the feeder in 1h as regular foragers.

### Training

We trained 20 foragers of each species in two visual conditioning protocols routinely used to assess bee learning and memory [26]. Based on preliminary observations showing that hornets in this context preferred sucrose solution to water (and never collected water), we used sucrose solution as positive reinforcement and water as negative reinforcement.

#### Differential conditioning

We trained individual hornets to associate a color A with a sucrose reward (positive conditioned stimulus CS+) and a color B with unrewarded water (negative conditioned stimulus CS-) for 10 consecutive trials [13]. In differential conditioning, the conditioned stimuli are unambiguously associated with an unconditioned stimulus or with its absence. This protocol was used to evaluate the learning ability of hornets.

#### Reversal conditioning

Immediately after differential learning, we trained the same hornets to learn the opposite association, so that the sucrose reward (CS+) was paired to color B and water (CS-) to color A during 10 additional trials [14]. In reversal conditioning, there is a transient ambiguity of stimulus outcome that needs to be overcome by the insect. This protocol was thus used to evaluate cognitive flexibility of hornets.

At each trial of each learning protocol, CS+ and CS-were randomly assigned in the arms of the Y-maze. Hornets were free to come in the Y-maze when motivated, which means that the inter-trial interval was not controlled (n=800 visits, mean ± SE: 7.35 ± 6.71 min, range: 1 to 116 min). We cleaned Y-maze with 70% ethanol after each trial to remove any potential chemical marks. For each hornet, we computed a learning score for differential learning and reversal learning by summing its choices during the 10 trials [27] (0: only CS-, 5: random, 10: only CS+). For each protocol, we considered that individuals that succeeded in the three last trials were “learners”.

### Memory retention

We tested short-term memory (STM) 1h after the last trial of reversal learning. This analysis was conducted only for learners. The trained hornets were allowed to re-enter the maze and choose between the two visual stimuli without any sucrose or water. We considered that the hornet memorized the association when it took the arm colored as CS+ and antennated the empty feeding hole. Opportunistically, when hornets returned at the maze entrance 24h after the STM test, we ran a similar test to assess long-term memory (LTM).

### Morphometry

To control for a potential influence of body size on learning and memory performances (as reported in other hymenopterans [28–30]), we froze killed the conditioned hornets and made morphological measurements with Toupview software coupled to a Nikon SMZ 745T dissecting microscope (objective x0.67) with a Toupcam camera model U3CMOS.

We measured head width as a proxy of body size [28]. Note that 4 out of the 20 *V. crabro* were physically damaged and removed from theses analyses.

### Statistics

We analysed the data in R 4.0.4. From the raw data (available in Dataset S1 and S2) we extracted the first choice of each hornet at each trial (CS+: 1, CS-:0) of differential and reversal conditioning. We then tested the influence of species (*V. crabro* or *V. velutina*) and trials (1-10) on first choice (CS+), using a generalized linear mixed model (GLMM; R package *lme4* [31]), with binomial error structure and identity as random factor, followed by an ANOVA (R package *car* [32]). For each conditioning protocol, we compared the number of learners in the two species using a Chisquare test with a continuity correction (R function *chisq*.*test*). We analysed the effect of head width on learning scores (0-10) using a linear model (R function *lm*).

## Results

We first assessed the learning performances of hornets in a differential learning task in which one colour was rewarded and the other was not (Figure 2A). The percentage of individuals that correctly chose the reward increased with experience (Binomial GLMM, trial: X^2^=43.53, df=9, p<0.001) and this was similar in the two species (Binomial GLMM, species: X^2^=2.26, df=1, p=0.13). The proportion of learners was identical (85%, 17/20 hornets) in *V. velutina* and *V. crabro*.

**Figure 2:**
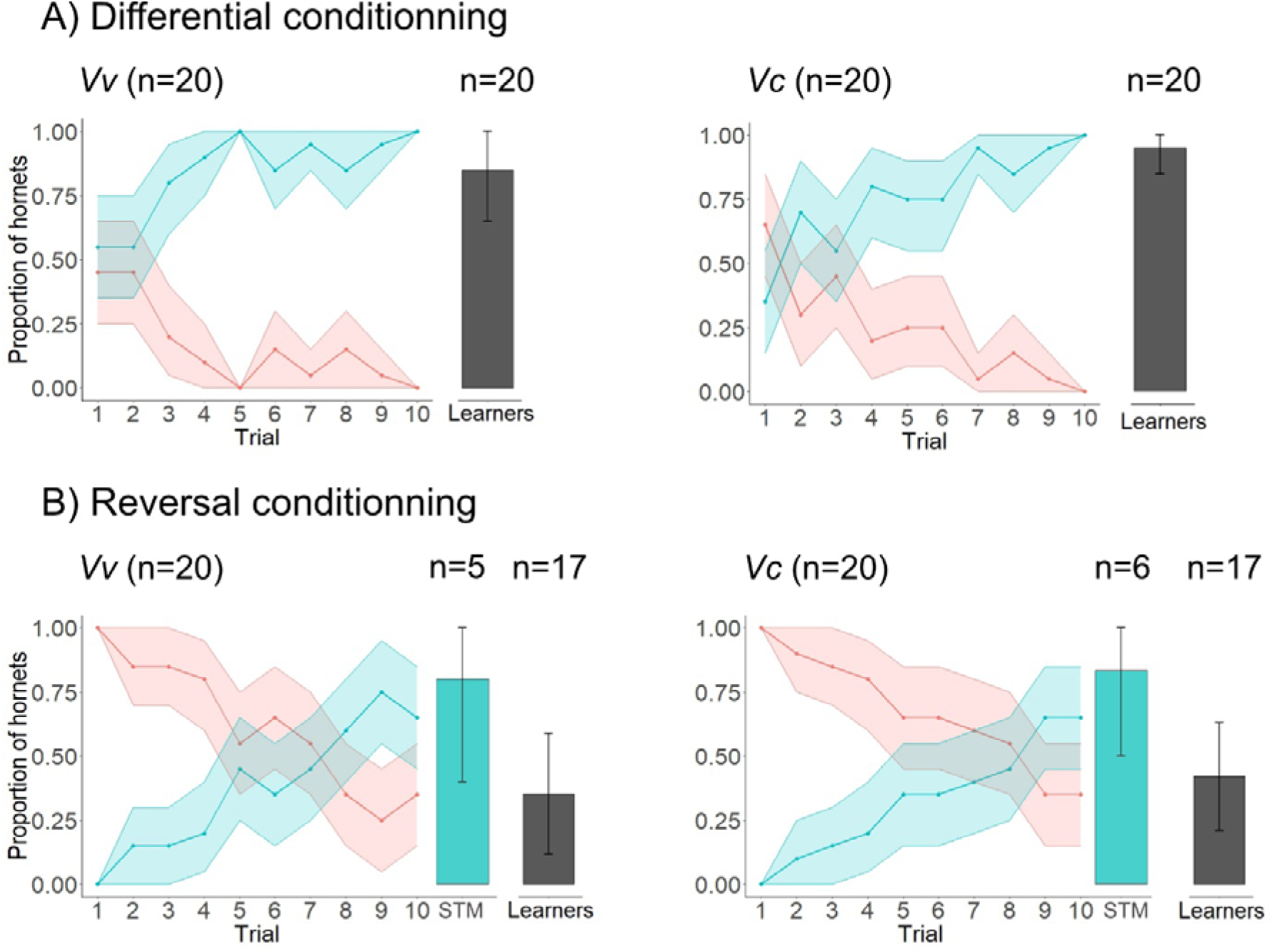
Learning curves for differential (A) and reversal (B) conditioning of *V. velutina* (Vv) and *V. crabro* (Vc). Red: responses to CS+. Blue: Responses to CS-. Learners: percentage of learners. STM: Short-term memory test. Error bars are 95% confidence intervals.

We then measured the cognitive flexibility of the learners in a reversal learning task where the previously learnt reward-colour associations were inversed (Figure 2B). Here again the proportion of hornets that chose the rewarded stimulus increased with experience (Binomial GLMM, trial: X^2^=0.53, df=9, p<0.001) and was similar in the two species (Binomial GLMM, species: X^2^=0.59, df=1, p=0.4). The proportion of learners was comparable in *V. velutina* (29.4%, 5/17 hornets) and *V. crabro* (35.2%, 6/17 hornets) (X^2^<0.001, df=1, p=1), but much lower than for differential learning, indicating that overcoming the transient ambiguity of the stimuli following the reversal of contingencies was complex.

In both species, most of the learners developed a short-term memory of the colour-reward association when tested 1h after the reversal learning phase (*Vv:* 80%, n=4/5; *Vc:* 83%, n=5/6). We also tested a subset of these learners for long-term memory 24h later. Four out the 11 individuals made successful choices (*Vv*: 1/5; *Vc*: 3/6), indicating that hornets can develop long-term memories of food-colour associations after at least ten trials (Supplementary Table 2). Our sample size was too small to test for statistical differences between species.

Head width varied by 13.46% (mean+SE: 3.37± 0.13 mm, range: 3.12-3.54 mm, n= 20) among *V. velutina* and 22.22% (3.86 ± 0.19 mm, range: 3.33-4.07, N = 20) among *V. crabro*. This had no influence on the learning scores (LM differential learning: *Vv:* X^2^=9.8, df=1, p=0.08, *Vc:* X^2^=4.73, df=1, p=0.19; LM reversal learning: *Vv:* X^2^=2, df=1, p=0.37; *Vc:* X^2^=4.32, df=1, p=0.33).

## Discussion

We adapted appetitive conditioning protocols used in bee research to explore and compare flexible visual cognition in nectar foraging hornets. Hornets of the two species learned equally well the visual-colour associations in differential and reversal conditioning, and remembered these associations for at least 24h. This suggests these forms of visual learning and cognitive flexibility are widespread in flower foraging animals.

Wasps are known to use visual cues in navigation [17,20,33,34] and communication [18,19,35]. However the importance of visual learning and memory in flower selection is less clear [21]. Here we showed that nectar foraging hornets exhibit cognitive performances comparable to those of bees tested in similar conditions [36,37]. Just like nectar foraging bees, *V. velutina* and *V. crabro* foragers can learn to associate colors with sucrose rewards and store these associations in memory for at least 24h. This memory of colour-reward association likely supports flower constancy, a behaviour recently described in hornets [38] and known to improve foraging success in bees [22]. Importantly, the hornets were also capable to replace the learnt associations by new ones, indicating that they can adjust their flower preferences to natural fluctuations of flower reward values through time, over the course of their foraging career. Such behavioural flexibility at the individual level may be critical for foragers to adapt nutrient collection to changing colony needs, depending on variations in colony composition (e.g. adult to larvae ratio) or external conditions (e.g. ambient temperature) [39].

The fact that we have detected no difference in the performances of *V. crabro* and *V. velutina* suggests that the cognitive traits we studied are basic abilities across nectar foraging species, irrespective of difference in morphology (body size) and invasion history between species. Y maze conditioning is a simple yet powerful approach for further analyses of the cognitive capacities of insect pollinators. Future studies using this approach could explore more elaborated forms of visual learning described in such non-elemental associative learnings [40] or bimodal visual-olfactory learning [41]. In the case of *V. velutina*, a detailed understanding of their cognitive abilities can help better predict their spreading dynamics (e.g. in relation to food type and abundance) or develop new tools for biocontrol (e.g. hornet traps exploiting learning and memory) of invasive populations in Europe and Western Asia. Ultimately, comparing cognitive abilities across species using standard, replicable tests, is critical to understand the ecological drivers of the evolution of cognitive traits.

## Acknowledgements

We thank Stephane Kraus helping producing Figures 1A-B.

## Contribution statement

MLa, DT and MLi designed the study. EN and CL built the Y-maze. MLa performed the experiments and wrote the first draft. MDR and NR participated to the experiments. All authors revised the manuscript.

## Supplementary materials

**Table S1:**
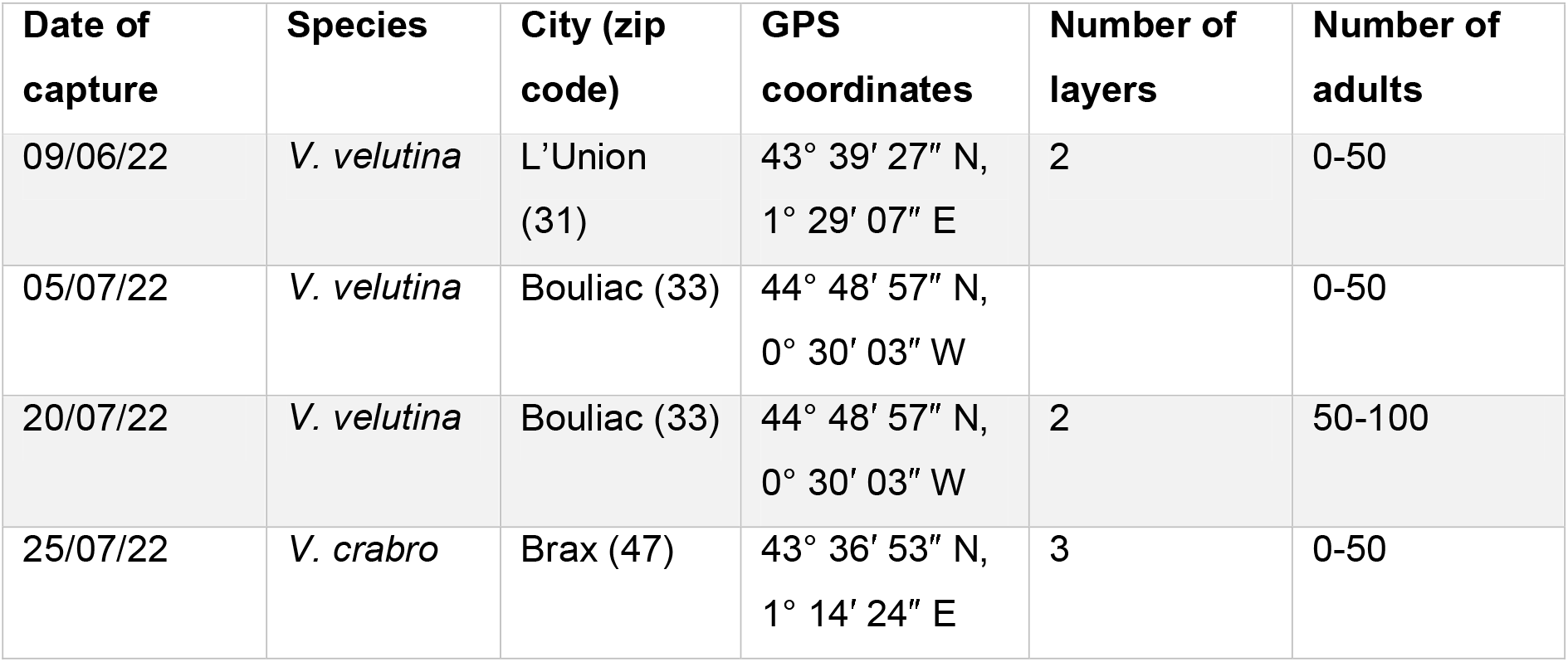
Date, origin of hornet nests and their composition

| Date of capture | Species | City (zip code) | GPS coordinates | Number of layers | Number of adults |
| --- | --- | --- | --- | --- | --- |
| 09/06/22 | *V. velutina* | L'Union (31) | 43° 39' 27" N, 1° 29' 07" E | 2 | 0-50 |
| 05/07/22 | *V. velutina* | Bouliac (33) | 44° 48' 57" N, 0° 30' 03" W |  | 0-50 |
| 20/07/22 | *V. velutina* | Bouliac (33) | 44° 48' 57" N, 0° 30' 03" W | 2 | 50-100 |
| 25/07/22 | *V. crabro* | Brax (47) | 43° 36' 53" N, 1° 14' 24" E | 3 | 0-50 |

**Table S2:** Summary of cognitive scores and memory retention results for each hornet.

| ID | Species<br>*V. velutina*<br>(Vv) or *V.*<br>*crabro*<br>(Vc) | Cognitive score<br>/10 |  | Learner<br>Yes: 3 last trials are<br>succeeded |  | Short<br>term<br>memory | Long<br>term<br>memory |
| --- | --- | --- | --- | --- | --- | --- | --- |
|  |  | Differential | Reversal | Differential<br>On all tested<br>individuals | Reversal<br>On learners<br>from<br>differential | Yes if<br>succeeded | Yes if<br>succeeded |
| 1 | Vv | 9 | 4 | Yes | No | Yes |  |
| 2 | Vv | 9 | 0 | Yes | No | No |  |
| 3 | Vv | 9 | 4 | Yes | No | Yes | Yes |
| 4 | Vv | 9 | 3 | Yes | No | Yes | Yes |
| 5 | Vv | 9 | 3 | Yes | Yes | yes |  |
| 6 | Vv | 10 | 3 | Yes | Yes | yes |  |
| 7 | Vv | 9 | 3 | Yes | No | Yes |  |
| 8 | Vv | 10 | 4 | Yes | No | Yes |  |
| 9 | Vv | 10 | 3 | Yes | No | Yes | Yes |
| 10 | Vv | 10 | 4 | Yes | Yes | yes |  |
| 11 | Vv | 9 | 5 | Yes | Yes | yes | Yes |
| 12 | Vv | 8 | 4 | Yes | No | Yes | No |
| 13 | Vv | 10 | 4 | Yes | No | Yes |  |
| 14 | Vv | 5 | 8 | No |  | yes |  |
| 15 | Vv | 5 | 3 | No |  | No |  |
| 16 | Vv | 7 | 6 | Yes | Yes | no |  |
| 17 | Vv | 4 | 4 | No |  | Yes | Yes |
| 18 | Vv | 8 | 3 | Yes | No | Yes | Yes |
| 19 | Vv | 9 | 3 | Yes | No | No |  |
| 20 | Vv | 9 | 3 | Yes | No | Yes |  |
| 21 | Vc | 5 | 0 | Yes | No | No | No |
| 22 | Vc | 8 | 4 | Yes | Yes | yes | Yes |
| 23 | Vc | 10 | 6 | Yes | Yes | yes | Yes |
| 24 | Vc | 7 | 6 | Yes | Yes | yes | No |
| 25 | Vc | 6 | 0 | Yes | No | yes | Yes |
| 26 | Vc | 9 | 0 | Yes | No | No | No |
| 27 | Vc | 7 | 4 | Yes | No | Yes | Yes |
| 28 | Vc | 9 | 1 | Yes | No | Yes | No |
| 29 | Vc | 4 | 8 | No |  | yes |  |
| 30 | Vc | 8 | 5 | No |  | Yes |  |
| 31 | Vc | 7 | 2 | No |  | Yes | Yes |
| 32 | Vc | 10 | 3 | Yes | Yes | no | Yes |
| **33** | *Vc* | 6 | 4 | Yes | No | Yes | Yes |
| **34** | *Vc* | 7 | 4 | Yes | Yes | yes | Yes |
| **35** | *Vc* | 9 | 5 | Yes | No | Yes | no |
| **36** | *Vc* | 10 | 2 | Yes | No | No | No |
| **37** | *Vc* | 8 | 3 | Yes | No | No | No |
| **38** | *Vc* | 6 | 4 | Yes | Yes | yes | No |
| **39** | *Vc* | 8 | 1 | Yes | No | Yes | No |
| **40** | *Vc* | 9 | 3 | Yes | No | Yes | Yes |

